# Isolated individuals and groups show opposite preferences toward humidity

**DOI:** 10.1101/398651

**Authors:** Mariano Calvo Martín, Stamatios C. Nicolis, Isaac Planas-Sitjà, Jean-Christophe de Biseau, Jean-Louis Deneubourg

## Abstract

Cockroaches, like most social arthropods, are led to choose collectively among different alternative resting places. These decisions are modulated by different factors, such as environmental conditions (temperature, relative humidity) and sociality (groups size, nature of communications). The aim of this study is to establish the interplay between environmental conditions and the modulation of the interactions between individuals within a group leading to an inversion of preferences. We show that the preferences of isolated cockroaches and groups of 16 individuals, on the selection of the relative humidity of a shelter are inversed and shed light on the mechanisms involved. We suggest that the relative humidity has a multi-level influence on cockroaches, manifested as an attractant effect at the individual level and as a negative effect at the group level, modulating the interactions.

## Introduction

Social animals are usually confronted to collectively choose the most suitable resource or resting place among several options in order to maintain social benefits. These collective choices are modulated by physiological (e.g. starvation, desiccation) and environmental (e.g. temperature, relative humidity) [Tremblay and Gries, 2006] factors; as well as by sociality (e.g. nature of communications, group size) [Chikao and Keiko, 1985, Krafft et al., 1986, Zahiri and Rau, 1998].

Many studies [van den Bos et al., 2013, King and Cowlishaw, 2007, Krause and Ruxton, 2002, Sumpter et al., 2008], have shown that the presence of conspecifics is able to amplify individual preferences through social interactions (interattractions between individuals), potentially leading to a better discrimination between resources quality [Canonge et al., 2009, Dussutour et al., 2005, Franks et al., 2002]. This classical approach of collective decision-making often underestimates however individual complexity and, in particular, the modulation of the interactions induced by the environment [Dambach and Goehlen, 1999, Hassanali et al., 1989]. It has been reported that individuals within groups show different preferences to those of isolated individuals (e.g. bark beetles (*Scolytinae*) attacking trees [Kausrud et al., 2011, Raffa and Berryman, 1983] or shelter selection in spiny lobster [Eggleston and Lipcius, 1992]) but the origin of the differences seems to be mainly related to crowding effects. A recent study showed a crowding-independent inversion of odour preferences between isolated individuals and groups [Laurent Salazar et al., 2017]. This type of phenomena, rather than being marginal should be at work in other situations involving the action of environmental factors on social species.

Most arthropods are highly sensitive to water losses and possess physiological (dryer feces, cuticular hydrocar-bon) and behavioural (hygrotaxis, clustering) mechanisms to counterbalance them [Edney, 1951, Gibbs et al., 1997, Leather et al., 1995, Reynolds and Bellward, 1989, Vanthournout et al., 2016, Yoder and Grojean, 1997]. Here, we test the preference between a dry shelter (DS) with a relative humidity (RH) of 40% and a wet shelter (WS) (RH 90%) offered to isolated individuals and groups of the American cockroach *Periplaneta american*a. The hypothesis put forward in this paper is that, depending on their physiological state, isolated individuals would prefer WS to avoid water losses [Bell and Adiyodi, 1982,Doi and Toh, 1992]. On the contrary, individuals in groups should prefer DS as high levels of humidity modulate negatively their interattractions [Dambach and Goehlen, 1999, Hassanali et al., 1989], which play an essential role in aggregation and collective decision making [Lihoreau et al., 2012].

## Results and discussion

The proportion of sheltered population over time (Figure 1a & Table 1) highlights the propensity of cockroaches to select dark resting places [Bell et al., 2007, Sempo et al., 2009]. At the end of the experiment, the mean sheltered proportion of isolated individuals (0.74) and groups (0.63) showed no statistical difference (*χ*^2^_1_ test _(N=518)_ =2.9; P=0.09).

**Table 1.** Different observations of the isolated and grouped trials and their respective statistic test.

|  | Observations | Both | Wet | Dry | Binomial test | Permutation test | W statistic (P) |
| --- | --- | --- | --- | --- | --- | --- | --- |
| Isolated<br>(N=54) | N° wins (proportion) | 40 (0.74) | 26 (0.48) | 14 (0.26) | 0.0403 * |  |  |
|  | N° entries | 283 | 170 | 113 |  | 0.01** |  |
|  | Mean N° entries (±SD) | 5.155 (±4.87) | 3.14 (±2.89) | 2.09 (±2.74) |  |  |  |
|  | N° exits | 243 | 144 | 99 |  | 0.036** |  |
|  | Mean N° exits (±SD) | 4.5 (±4.73) | 2.66 (±2.67) | 1.8 (±2.73) |  |  |  |
|  | Total Time | 280371.2 s | 173213 s | 107158.2 s |  |  | 1730.5 (0.04)*** |
|  | Mean total time (±SD) | 5192 s (±3768.1 s) | 3207.6 s (±3725 s) | 1984.4 s (±3401.2 s) |  |  |  |
|  | Mean visit time (±SD) | 990.7 (± 24165.5) | 962.2 s (±2369.1 s) | 830.6 s (± 2277.8 s) |  |  | 13850 (0.003)*** |
| Grouped<br>(N exp=29)<br>(N ind= 464) | N° Sheltered after 3 h (proportion) | 289 (0.62) | 113 (0.24) | 176 (0.37) | 0.0001**** | 0.0046** |  |
|  | Mean sheltered ind./trials | 10,04 (±4.37) | 3.98 (±3.91) | 6.06 (±4.21) |  |  |  |
|  | N° wins | 28 | 8 | 20 | 0.035**** |  |  |
|  | N° entries | 1840 | 988 | 852 |  | 0.11** |  |
|  | Mean N° entries tirals (±SD) | 63,44 (± 29.93) | 34.06 (±18.99) | 29.37 (±17.68) |  |  |  |
|  | N° exits | 1551 | 875 | 676 |  | 0.0017** |  |
|  | Mean N° exits tirals (±SD) | 53.48 (±28.66) | 30.17 (±17.25) | 23.31 (±15.81) |  |  |  |

\* One tail binomial test, H0: the probability of observation for the wet shelter is $> 0.5$ .
\*\*Permutation test (100000 repetitions): probability of observing the experimental results.
\*\*\*One tail Mann–Whitney–Wilcoxon test, H0: The observed distribution in the wet shelter is greater than in dry shelter.
\*\*\*\* Binomial test, H0: The probability of observation is 0.5 in both shelters.

**Figure 1.**
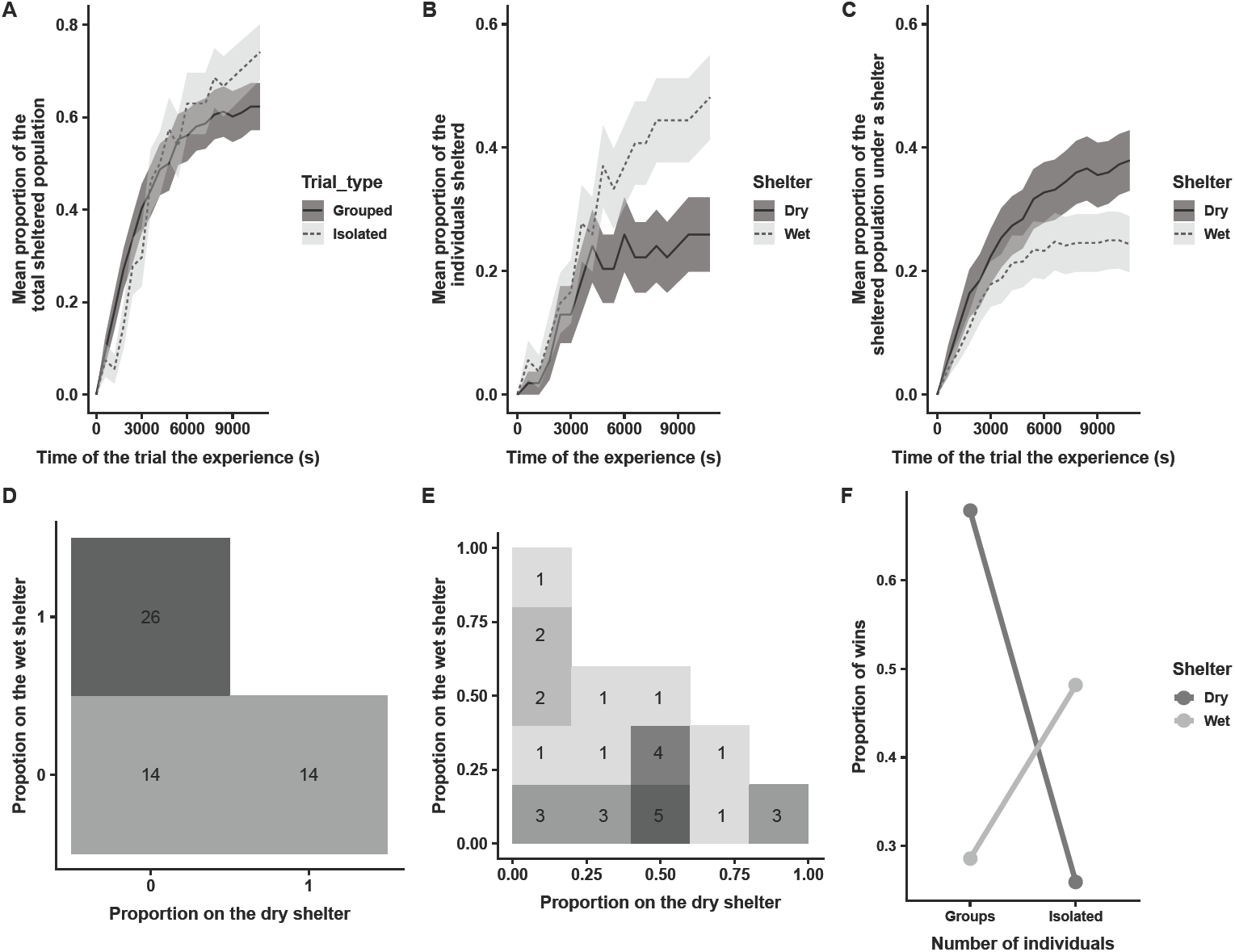
(A) Mean ± SE proportion of sheltered individuals over time for isolated (light grey) and groups (dark grey). (B) Mean ± SE proportion of the isolated individuals over time for the dry (dark grey) and the wet (light grey) shelters. (C) Mean ± SE proportion of individuals in groups over time for the dry (dark grey) and the wet (light grey) shelters. (D) Distribution of the presence/absence under the dry (dark grey) and wet (light grey) shelters for the isolated individuals’ trials. (E) Distribution of the sheltered population under the dry (dark grey) and wet (light grey) shelters for the groups trials. (F) Proportion trials with most of individuals choosing the dry (dark grey) and wet (light grey) shelter, at the end of the experiences, for the isolated individuals and the groups.

Isolated cockroaches (figure 1b; d) settled more frequently under WS than under DS (0.65 of the total sheltered individuals at the end of all experiments; one-tail binomial test P =0.04). This is in agreement with studies showing that isolated individuals actively search for a resting place with high humidity levels, allowing a reduction of water losses [Ramsay, 1935]. However, the situation is inverted for the cockroaches in groups which collectively select the resting place with lower RH level (0.61 of the total sheltered population; binomial test P =0.0001) (Figure 1c; e).

In groups, interattractions between individuals are at work and therefore the individual choices are not independent. Indeed, if the settlement and the shelter selection were the result of a non-social behaviour, the distribution of sheltered populations would follow a binomial distribution with a probability of 0.607 (mean proportions of sheltered individuals under DS at the end of the experiments) to be settled under DS. The variance of the population under the DS among the replicas (17.8) is significantly higher than the theoretical variance (binomial distribution, see material and methods; P*<*0.001). A permutation test (see material and methods) shows that the probability to settle under DS was significantly larger than the probability to settle under WS (P=0.034). Note that due to social interactions, this system can display multistationarity and therefore some trials end with the selection of WS (see figure S1). Even so (see figure 1f), summing all trials ending with most of individuals choosing WS (DS) shows clearly the inversion between isolated individuals and groups (*χ*^2^_1 (N=68)_ =8.743; P=0.003).

The proportion of entries (exits) under (from) a shelter is related to the probabilities of joining (leaving) both shelters (see material and methods) and to the time bouts spent outside (inside) this shelter. In the isolated trials the number of entries in WS was higher than DS (proportion=0.6; table1; permutation test P=0.01). For the groups, the number of entries was also higher, but no significant difference was observed (proportion=0.53, table 1; permutation test P=0.11). This proportion was constant regardless the number of individuals inside the shelters (See supplementary). As for the exits, the comparison of the time bouts spent under WS or DS showed that the probability of leaving WS is lower for the isolated individuals (table 1). For the groups, the exit proportion from WS was higher than 0.5 (0.56) despite the fact that its population was lower (table 1) and the permutation test showed that the exits from both shelters are statistically different (higher for WS, P=0.017).

All in all, the RH% has an attractive effect for the isolated individuals and groups, although, this attraction is weaker for the groups. For the isolated individuals the probability of leaving is lower for WS than for DS. The settlement under WS is thus more frequent for isolated individuals. This was not observed for the groups where the settlement under the DS was more frequent and more populated than under WS. In this case the probabilities of joining both shelters were similar but the probability of leaving DS was lower. It has been shown that individuals emit attractant volatile components [Säid et al., 2005, Tanaka and Daimon, 2018], which in the framework of our experiments with groups may have led to a decrease of the relative attractiveness of WS versus DS. On the other hand, [Dambach and Goehlen, 1999, Hassanali et al., 1989] showed that high levels of humidity decrease the interattractions which play an essential role in sheltering behaviour [Ame et al., 2006, Halloy et al., 2007, Jeanson and Deneubourg, 2007]. According to our hypothesis, for identical population sizes under WS or DS, this decrease of interactions leads to a higher probability of leaving from WS. Stated differently, as the population under DS increases, the probability of leaving decreases and the aggregate under this shelter compensates water losses caused by dryness [Vanthournout et al., 2016, Yoder and Grojean, 1997]. In fact, the probabilities associated to the sheltering behaviour depend not only on the different humidity levels but also on the already sheltered population. This modulation coupled with the competition generated by the presence of two shelters, are the two ingredients at the basis of the patterns observed.

Low interest for high humidity levels by the groups might be related to a prophylactic strategy to reduce pathogen transmission. Indeed, higher humidity levels increase the speed of fungal development [Doberski, 1981, Mishra et al., 2015] and the mortality rate [Hernandez-Ramirez et al., 2008] while it has been suggested that low humidity levels reduce horizontal transmissions of pathogens [Quesada-Moraga et al., 2004]. Furthermore, the composition of the cuticular hydrocarbons should play an essential role in the aggregation process, as they are linked to the resistance to desiccation [Gibbs et al., 1997, Säid et al., 2005], and constitute a first barrier against pathogen [Pedrini et al., 2013]. Tackling quantitatively these bio-physico-chemical mechanisms and their relation to collective behaviour in social species would help to disentangle the distal causes of the inversion of the preferences between isolated individuals and individuals in groups observed in this study.

## Materials and methods

### Biological model

*Periplaneta americana* (L.) (Dictyoptera: Blattidae) is a domiciliary cockroach that forms aggregates during daylight hours in dark places and is active during night-time. The cockroaches used in this study measured from 35-50 mm in length and were issued from strains reared in breeding facilities (five Plexiglas vivaria of 80×40×100cm) of the Université libre de Bruxelles. Each vivarium contained about 1000 individuals of both sexes and of all developmental stages and we provided dog pellets and water twice a week. The rearing room was maintained at 25 *±* 1 *°* C and 40% RH under a 12:12 h light/dark cycle.

### Experimental setup

Experiments were carried out only with male adults of *P. americana* without external damage, to exclude any behavioural variation linked to ovarian cycle. The experimental set-up (figure S3) was a circular arena, covered with a paper layer (120 g/m^2^), surrounded by a polyethylene ring (diameter: 100 cm, height: 20 cm) with a light source (5 M Ustellar Dimmable Kit Ruban Led, 2835 SMD Led, White cold 6000 K) placed above the set-up and providing a homogeneously light intensity of 415 lux at ground level. To avoid any visual cue, the arena was inside a white box of 105×105×130 cm (WxLxH). Two shelters (one dry, one wet see below) made of transparent Plexiglas pipes (H: 30cm; D: 15cm) covered with red-coloured filter film (Rosco E-color 19: fire). Their upper side was covered with a black carton. The light intensity inside the shelter is of 22 lux. A transparent ceiling was used at 2,5 cm to reduce the volume of the shelter. The centre of each shelter was located 23 cm from the edge of the arena.

Shelters had two symmetrically opposed entrance of 2×1,5 cm (WxH) aligned to the centre of the arena. To control humidity inside the shelter, a hole of 11 cm covered with a plastic grid was made in the floor of each shelter. The floor of the arena was covered with a paper layer, with two openings at the place of the shelter. This paper was changed and the shelters, as well as the floor under each shelter, were cleaned after every trial, to avoid any chemical marking. Each shelter was large enough to accommodate at least 16 cockroaches.

### Humidity control

Cockroaches are photophobic and both shelters therefore are perceived as resting sites during the diurnal phase [Bell and Adiyodi, 1982]. Moreover, cockroaches are sensitive to the “dryness” or “wetness” of the environment, which is reflected by the saturation deficit (SD) [Tichy and Kallina, 2013]. It depends on the temperature as well as the relative humidity. Since all the trials were conducted at 25 *±*2*°*C, the SD was only dependent on the relative humidity (RH). The “dry” shelter had the same RH as the experimental room, which was maintained at 25 *±*2*°*C and a RH of 35 - 50 %. The wet shelter had a RH of 92 *±*5%, generated by adding 40 ml of tap water [Dambach and Goehlen, 1999] on a petri dish, 10 cm beneath the plastic grid. The room conditions were measured with a multi-function climate measuring instrument (Testo 435 coupled to a temperature and humidity probe). Humidity and temperatures under the shelter were measured with humidity and temperature sensors (DHT22) beneath the plastic grid from the floor of the shelter.

### Data recording

To allow the detection of the animals under the shelters, the cockroaches were tagged with an ArUco tag [Romero-Ramirez et al., 2018]. On their thorax with Latex (Winsor & Newton) sheltered individuals were detected by a video camera located on the upper side of the shelter.

### Behavioural assay

Groups of 16 cockroaches were kept in total darkness for 48 h in Plexiglas ® boxes (36 x 24 x 14 cm) containing a cardboard shelter, humidified cotton wool and *ad libitum* food.

A single male or a group 16 individual was introduced at the centre of the arena and the animals were free to explore it and visit the shelters. Each trial lasted 3 h. This procedure was repeated 54 times for isolated individuals and 29 times for groups. For each shelter, we measured the number of entries and exits, the number of sheltered individuals as function of time and the time bouts (only for the isolated trials).

### Data and statistical analysis

The recording of the experiment was analysed using Solomon coder https://solomoncoder.com/). To test preferences on the selection of a resting place depending on the RH. We use binomial tests on the number of total individuals under each shelter at the end of the experiences, for the isolated individuals (H0: the probability of a cockroach to choose the wet shelter at the end of the trial is greater than 0.5) and for the groups (H0: the total shelter population at the ends of all trial are the same for the wet then for the dry). *χ*^2^ test was used to test differences between isolated and groups for the total sheltered populations and the most populated shelter. For groups of 16 cockroaches, we tested whether the observed distribution of sheltered individuals follow a binomial distribution. For this purpose, we compute the theorical mean and variance of the population under DS using a simulation (100,000 realisations) of the observed total sheltered population using as probability of sheltering under DS the proportion of the total sheltered population of DS at the end of all trials. Moreover, a permutation test (100,000 x 29 permutations) was used to test if the number of individuals and the numbers of entries(exits) under each shelter differ from a of a distribution with an equal probability to settle under each shelter [Good, 2005]. Regarding the isolated individuals, we fallowed the same procedure (1000000 x 54 permutations) and a Wilcoxon-/Mann-Whitney test was used compare the total sheltering time and the time bouts spent under each shelter. The significance of statistical tests was fixed to *α* = 0.05. The probabilities of joining(leaving) the shelters were calculated as follows:

• the relative probabilities of joining the dry(wet) shelter expressed as

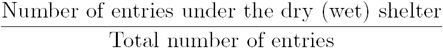

and

• The relative probability of leaving the dry(wet) shelter that can be written as

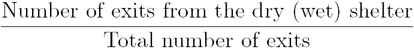

Note that for the isolated individuals we also calculated the probability of leaving per time unit Which reads

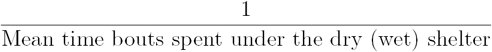

Finally, for the groups, we performed a linear regression to test the influence of the total number of sheltered individuals on the proportions of entries for the wet shelter.

Data analyses and permutations were performed using R software (R Core Team 2018, R Foundation for Statistical computing, https://www.r-project.org/) and Python (Python Software Foundation. Python Language Reference, version 2. 7. 15 at).

## Author contribution

M.C.M. participated in the conception of the study, carried out the experiments and participated in the data analysis and draft the manuscript. S.C.N. participated in the conception of the study, the data analysis and helped draft the manuscript. I.P.-S. participated on the graphical analysis and helped draft the manuscript. J.C.d.B co-supervised the study. J.L.D. participated in the conception of the study, the data analysis and helped draft the manuscript and co supervised the study.

## Supplementary

**Figure S1.**
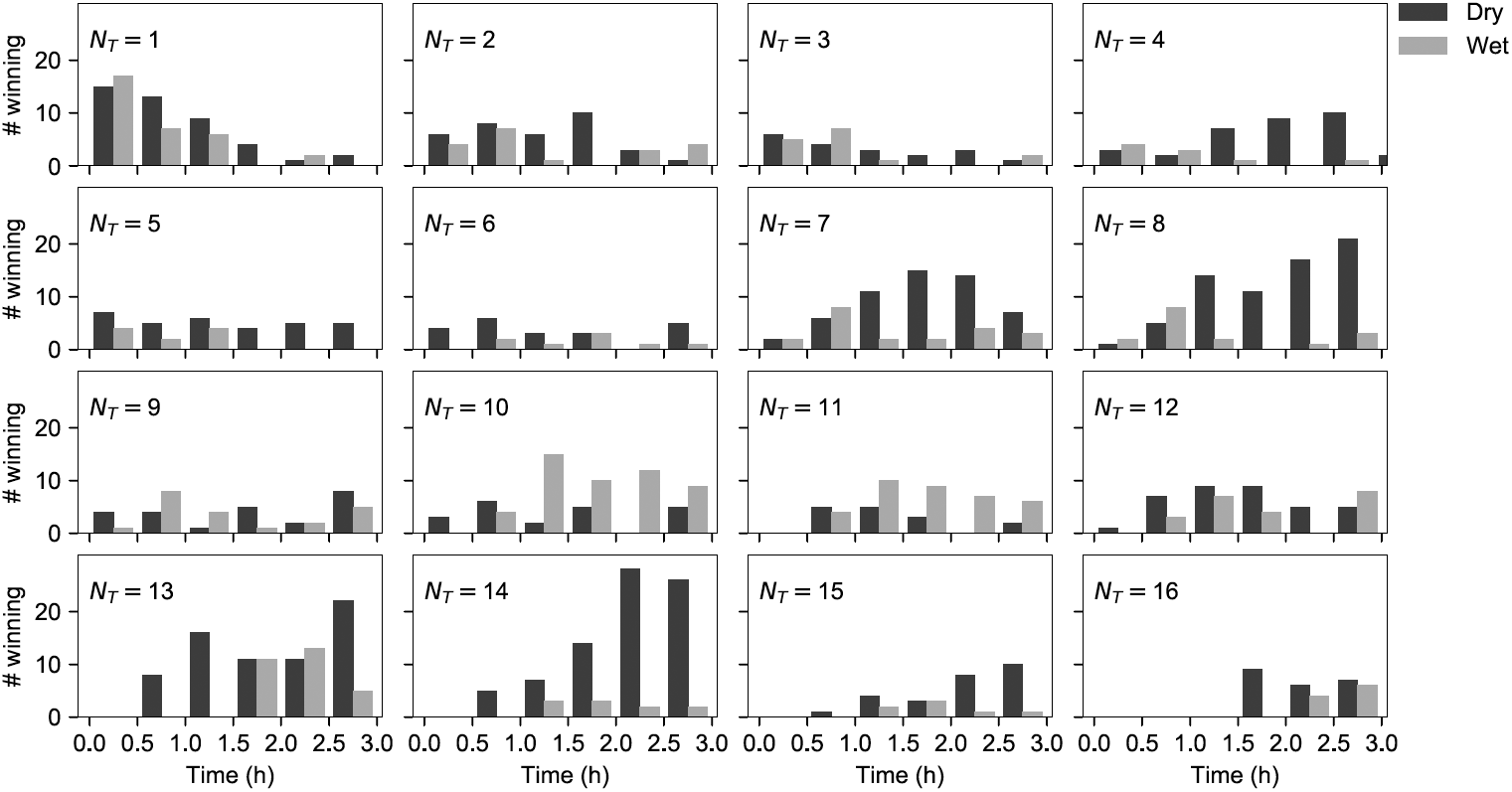
Number of times the number of individuals under DS (dark grey) or under WS (light grey) is larger than the number of individuals in the other shelter as a function of time, for each total sheltered population.

**Figure S2.**
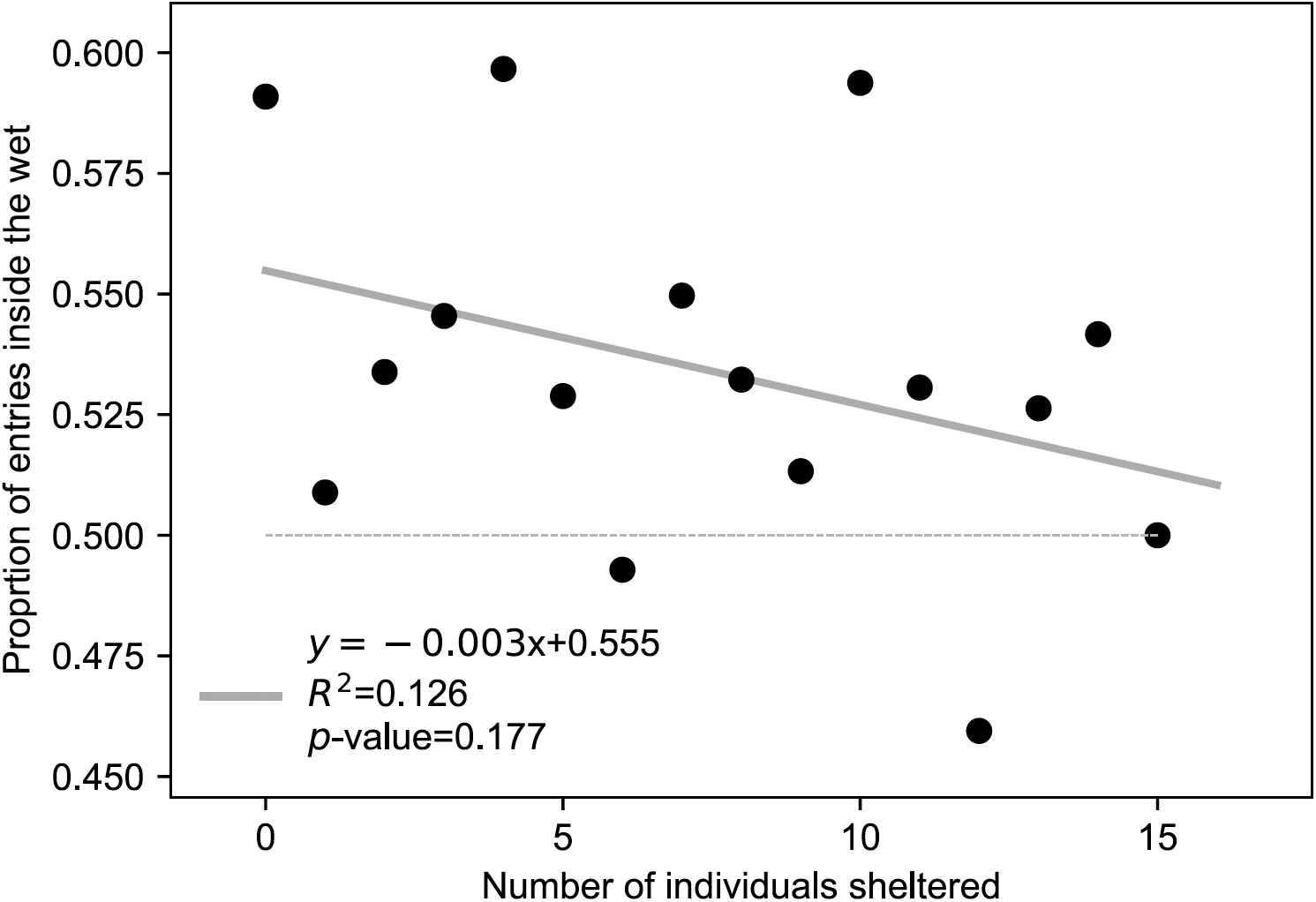
Proportion of entries in WS as a function of the total sheltered population along with linear regression.

**Figure S3.**
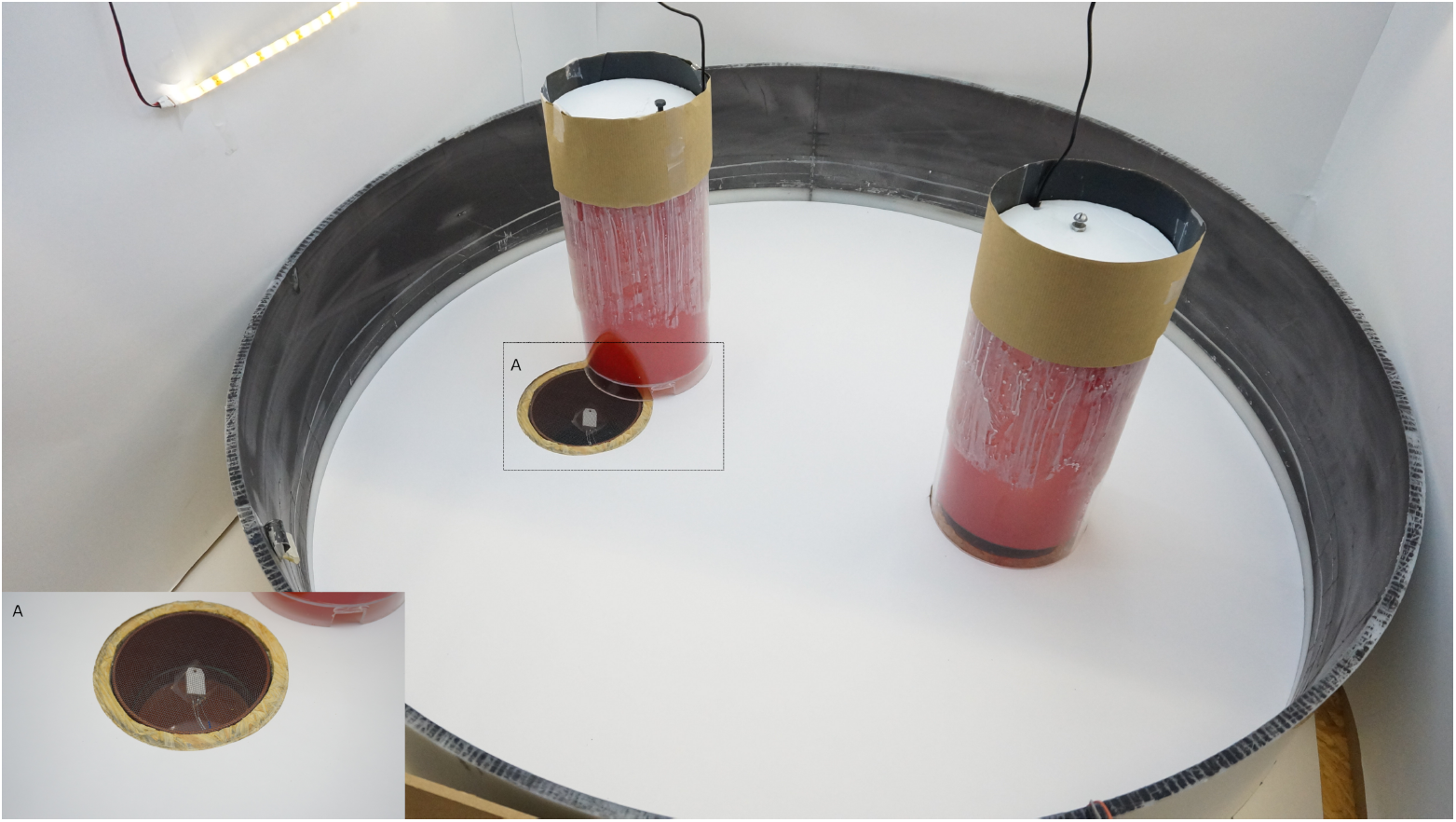
Experimental set-up, A) Zoom on the floor of the shelter.

**Figure S4.**
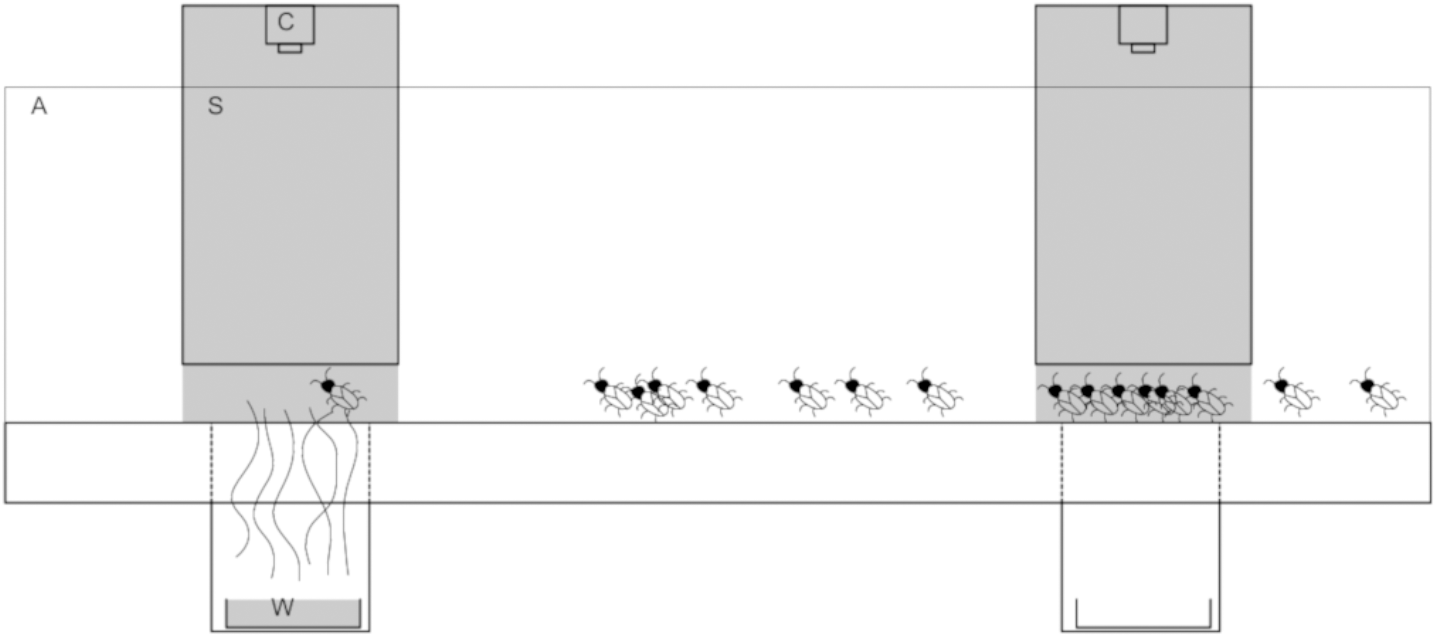
Scheme of the experimental set-up. A) Circular arena; C) Camera; S) Shelter; W) Source of humidity (water)

